# Cytokine and Leukocyte Profiling Reveal Pro-Inflammatory and Autoimmune Features in Frontotemporal Dementia Patients

**DOI:** 10.1101/049791

**Authors:** Philipp A. Jaeger, Trisha M. Stan, Eva Czirr, Markus Britschgi, Daniela Berdnik, Ruo-Pan Huang, Bradley F. Boeve, Adam L. Boxer, NiCole Finch, Gabriela K. Fragiadakis, Neill R. Graff-Radford, Ruochun Huang, Hudson Johns, Anna Karydas, David S. Knopman, Michael Leipold, Holden Maecker, Zachary Miller, Ronald C. Petersen, Rosa Rademakers, Chung-Huan Sun, Steve Younkin, Bruce L. Miller, Tony Wyss-Coray

## Abstract

The growing link between systemic environment and brain function opens the possibility that cellular communication and composition in blood are correlated with brain health. We tested this concept in frontotemporal dementia with novel, unbiased tools that measure hundreds of soluble signaling proteins or characterize the vast immune cell repertoire in blood. With these tools we discovered complementary abnormalities indicative of abnormal T cell populations and autoimmunity in frontotemporal dementia.

## INTRODUCTION

Frontotemporal Dementia (FTD) is a neurodegenerative disorder characterized by progressive atrophy of the frontal and temporal lobes and a leading cause of early onset-dementia (1, 2). There are three main forms of FTD: two language variants classified as progressive non-fluent aphasia or semantic-variant primary progressive aphasia (svPPA; also known as “Semantic Dementia”), as well as a behavioral variant (bvFTD) (1, 3). Clinically, svPPA presents with prominent deficits in the ability to understand word meaning, loss of empathy, and anterior temporal lobe atrophy (1). SvPPA has the highest clinicopathological specificity, presents with characteristic neuronal aggregates of TAR DNA binding protein 43 (TDP-43) (1) and is the least likely of the FTD variants to be genetically determined. Recently, a whole exome association study of sporadic FTD patients, found no evidence of specific genetic variations driving the disease (4). This suggests that external factors and multi-organ involvement may play a role in disease onset and progression (5), making it an ideal candidate to explore these complex systemic interactions.

We have previously reported the clinical observation that svPPA patients exhibit signs of increased autoimmune comorbidities and elevated levels of pro-inflammatory cytokine Tumor necrosis factor alpha (TNFα) in plasma (6), indicative of a strong neuro-immunological component to svPPA pathology. Sjögren *et al*. describe elevated TNFα levels in cerebrospinal fluid of pathologically uncharacterized FTD patients (7), suggesting that immune-modulatory therapies might prove beneficial in this condition. Additionally, in Paget Disease of Bone and Frontotemporal Dementia syndrome (another TDP-43 proteinopathy), Dec *et al*. report elevated TNFa levels in plasma (8). In a remarkable coincidence, the growth factor progranulin (GRN), which is mutated in a rare familial form of FTD, directly binds to TNFa receptors as a TNFa antagonist (9, 10) (although this has been disputed by Chen *et al*. (11)), is targeted by autoantibodies in various autoimmune disease (12), and modulates neuroinflammation and T-cell proliferation (13, 14). However, in svPPA it remains unclear how TNFα and autoimmunity interact and if other immune pathways are involved (**Fig. 1A**).

**Figure 1:**
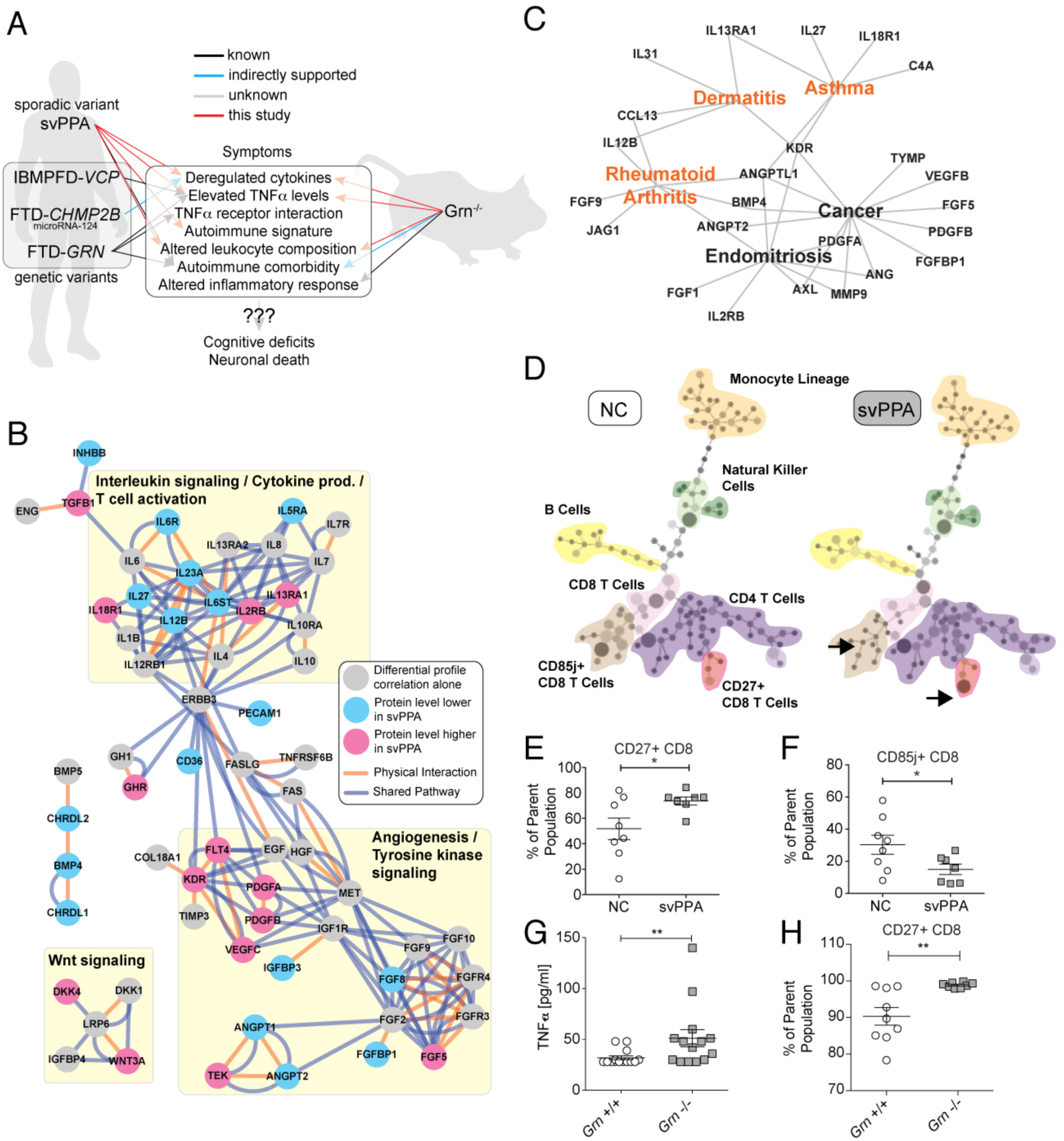
The role of systemic immune changes in Frontotemporal Dementia. **(A)** Overview of the finding in this study, integrating mouse model and human data. The big picture emerging is a widespread deregulation of inflammatory response, probably involving TNFα signaling, which in turn could promote the development of certain autoimmune conditions *via* altered leukocyte composition / activity. **(B)** Using an unbiased analysis of secreted plasma proteins, we identified numerous changes in cytokines and growth factors, implicating “Interleukin signaling” and “Angiogenesis” as potential target pathways (the Interleukin changes are most specific to svPPA). **(C)** Functional disease ontology analysis revealed shared features between svPPA and prominent autoimmune conditions. **(D-F)** We observed changes in leukocyte composition in svPPA patients through high-resolution immunophenotyping (NC: Non-demented controls). **(G)** In *Grn*^-/-^ mice, blood plasma TNFα levels were increased. **(H)** In the same mice, we also observed leukocyte changes similar to the ones found in svPPA patients.

## RESULTS

Here we present data towards answering these questions by analyzing patient and mouse blood samples, both for plasma protein and immune cell composition. First, we performed a targeted antibody array screen (15) of 593 secreted proteins involved in cellular signaling in plasma from 92 patients with svPPA and 83 cognitively normal controls (NC, **Table S1** and **Supplemental data file**). Unsupervised clustering indicated the presence of distinct disease related protein patterns (**Fig. S1, Extended Results and Discussion**). Gene ontology analysis showed that proteins, which are significantly different between svPPA and NC (FDR<0.05), are involved in cellular development, differentiation, and growth factor binding (**Table S2 and S3**). By utilizing a novel protein profile correlation-based approach, we were able to identify additional proteins that systematically changed their plasma levels in a parallel fashion, suggestive of related function (**Fig. S2, Extended Results and Discussion)**. A pathway-level analysis of the resulting set of plasma proteins indicated several functional clusters: “interleukin signaling / T-cell activation / cytokine production”, “angiogenesis”, and “Wnt signaling”. These findings are suggestive of a more widespread cytokine deregulation beyond TNFα alone, potentially involving leukocyte activation (**Fig. 1B**). To determine the specificity of these implicated pathways, we collected 47 additional samples from patients diagnosed with Alzheimer’s disease (AD, **Table S4**) and performed a similar plasma protein analysis (Jaeger et al., Molecular Neurodegeneration, in press). We found that out of 44 factors significantly changed in svPPA (FDR < 0.05), 17 factors were also changed in AD (FDR<0.05), while 27 factors were unique to svPPA (**Supplemental data file**). The unique factors were almost exclusively part of the “interleukin signaling / T-cell activation / cytokine production” pathway (**Fig. 1B** and **Supplemental data file**). After using Functional Disease Ontology annotations (16) to associate these plasma proteins with specific diseases, we noticed several autoimmune conditions (**Fig. 1C** and **Supplemental data file**). This led us to the hypothesis, that numerous cytokine-mediated changes generate an imbalance between pro-and anti-inflammatory activities in svPPA patients.

To follow this lead, we investigated the immunological changes that might meditate or be the result of such cytokine abnormalities. We performed a detailed immune phenotyping for 34 cell surface markers in peripheral blood monocytes (PBMCs) in an independent cohort of 8 NC and 7 svPPA patients using CyTOF (**Extended Results and Discussion, Table S5** and **S6, Fig. S3**) (17, 18). We observed altered populations of T cells (**Fig. 1D**). Specifically, significantly more CD3+ CD8+ T cells expressed CD27 (**Fig. 1E**) and fewer CD3+ CD8+ T cells expressed CD85j (**Fig. 1F**) in svPPA patients compared to NC. These results are consistent with the deregulation of IL-12/IL-23 signaling observed in the svPPA patients (**Fig. 1B**). These observations suggest that svPPA patients may have an immune system that may be conducive to developing (auto)-inflammatory diseases, thus providing a molecular explanation for clinical comorbidities (6).

No mouse model is currently available to investigate sporadic svPPA. However, we had previously reported that patients with familial FTD harboring mutations in *GRN*, a gene involved in immune regulation (19), exhibited elevated TNFa levels and autoimmune comorbidities (6). Thus, it would be intriguing if mice lacking this gene would mimic FTD-related immune abnormalities. Indeed, TNFα is elevated across these sporadic and genetic forms of FTD (6), as well as in *Grn^-/-^* mice (**Fig. 1I**), making immune system involvement likely. Also, *Grn^-/-^* mice have been shown to exhibit increased susceptibility to collagen-induced arthritis (9). By measuring basal cytokine levels in *Grn^-/-^* mice and littermate, wildtype controls using the Luminex platform, we observed widespread cytokine abnormalities, supporting previous findings of altered cytokine release following bacterial endotoxin stimulation (20). However, the detailed protein changes in mice differed from the human svPPA patients (**Supplemental data file, Extended Results and Discussion**). To determine if the mouse model shares the T cell phenotype observed in svPPA patients, we harvested PBMCs from *Grn^-/-^* and wildtype mice and used flow cytometry to measure populations of CD3+ CD8+ CD27+ cells. We found that significantly more CD3+ CD8+ T cells express CD27 in *Grn^-/-^* compared with wildtype mice (**Fig. 1J**), analogous to patients with svPPA (**Fig. 1C**).

## DISCUSSION

In summary, we report here multiple lines of evidence that svPPA patients exhibit widespread changes in blood cytokine levels, corroborating isolated cerebrospinal fluid findings from previous studies (21). The cytokine changes associated with changes in specific leukocyte subpopulations, suggestive of cellular immune abnormalities in the disease. TNFα is elevated in both sporadic and familial patients and mouse models of FTD, indicating that the immune involvement is significant to genetic and non-genetic forms of the disease. This notion is supported by specific changes in the immunephenotype of *Grn^-/-^* mice, mimicking findings form the svPPA patients. Intriguingly, another genetic variant of FTD that is mediated by CHMP2B mutations has recently been shown to cause changes in microRNA-124 (1, 2, 22), which in turn is a regulator of TNFα and IL6 activity (**Fig. 1A**) (1, 3, 23). Our findings thus suggest that different pathological manifestations of FTD subtypes appear to share a common immune phenotype that might be linked to abnormalities in the central nervous system and possibly neurodegeneration. In the light of these results and the high prevalence of FTD, further investigation of the optimal immune-modulatory therapeutic strategy appears both promising and timely.

## MATERIAL AND METHODS

### Participants

All participants underwent thorough and standardized history and physical exam. A total of 92 svPPA patients from the University of California San Francisco or Mayo Clinic Jacksonville were identified whose clinical features conformed to revised consensus diagnostic criteria for svPPA (6, 24) and had available serum and or whole blood for investigation (**Supplemental Table S1**). A total of 83 age, and gender matched healthy controls were obtained from a set of recruited controls from healthy aging programs at the University of California San Francisco or Mayo Clinic Jacksonville. Participants were considered to be healthy if they had normal neurological exam, MRI scans without clinically apparent strokes, without cognitive deficits on detailed neuropsychological testing, and without diagnosis of major psychiatric disease. Details about the AD comparison cohort are described elsewhere (Jaeger et al., under review). In short, samples from 47 AD patients and respective controls (**Supplemental Table S3**) were collected at the same centers as the svPPA and control samples and processed in parallel. Informed consent was obtained from human subjects according to the ethics committee guidelines at the respective clinical centers.

### Plasma Sample Preparation and Antibody-Microarray Incubation

The human plasma samples were platelet-and lipid-reduced by centrifugation. The plasma was then diluted and dialyzed (96 well Dispodialyzer/5kDa, Harvard Apparatus, Holliston, MA) into PBS (pH 6.5) at 4°C in multiple over-night steps to yield a maximally pure plasma protein fraction in an appropriate buffer for the biotinylation reaction. The plasma proteins were N-terminally biotinylated (EZ-Link NHS-Biotin, Thermo Scientific, Rockford, IL), unbound biotin removed by dialysis against PBS (pH 8.0) and the individual samples were incubated on blocked antibody arrays over-night at 4°C. After multiple washing steps antibody-bound protein was detected using Alexa Fluor 555 conjugated streptavidin (Invitrogen) on a GenePix Pro 4000B scanner (Molecular Devices, Sunnyvale, CA).

### Plasma Protein Antibody-Microarray Production

Plasma protein levels were measured using antibody-based protein microarrays. We used a custom-expanded, commercially available microarray with modified antibody content (custom L-Series, RayBiotech Inc., Norcross, GA) containing 474 antibodies against chemokines and cytokines printed in triplicates by the company, plus 17 control antibodies. Additionally, we produced a custom-made in-house array that contained a separate set of 119 antibodies against secreted signaling factors printed in quadruplicates, plus 10 control antibodies. A total of 593 antibodies against secreted signaling factors were measured. Subsequent quality control steps removed 11 antibodies with extremely low or no signal (see **Antibody-Microarray Data Preparation** below) yielding a total of 582 analyzed antibodies (**Supplemental Data File**). Some antibodies target the same protein multiple times (such as precursor/full-length/truncated forms, 14 proteins and 32 antibodies total; **Supplemental Data File**). The microarray production protocol was the following: antibodies of interest were selected based on their biological role as secreted signaling factors and the availability of ELISA-grade quality batches to ensure likely detection of the epitope in liquid solution. The arrays were printed onto SuperEpoxy glass slides (ArrayIt, Sunnyvale, CA) using a custom-built robotic microarrayer fitted with sixteen SMP4B pins (Arrayit). After drying the slides overnight they were vacuum-sealed and stored at −20°C until use.

### Plasma Sample Preparation and Antibody-Microarray Incubation

The human plasma samples were platelet-and lipid-reduced by centrifugation. The plasma was then diluted and dialyzed (96 well Dispodialyzer/5kDa, Harvard Apparatus, Holliston, MA) into PBS (pH 6.5) at 4°C in multiple over-night steps to yield a maximally pure plasma protein fraction in an appropriate buffer for the biotinylation reaction. The plasma proteins were N-terminally biotinylated (EZ-Link NHS-Biotin, Thermo Scientific, Rockford, IL), unbound biotin removed by dialysis against PBS (pH 8.0) and the individual samples were incubated on blocked antibody arrays over-night at 4°C. After multiple washing steps antibody-bound protein was detected using Alexa Fluor 555 conjugated streptavidin (Invitrogen) on a GenePix Pro 4000B scanner (Molecular Devices, Sunnyvale, CA).

### Antibody-Microarray Data Preparation

All data normalization and analysis was performed with custom code in Matlab (Mathworks, Natick, MA). Individual array spots were median background subtracted locally and spots that were not at least 10% more intense than median local background were defined as “non-detectable”. Antibodies with more than 55% non-detectable spots were removed from the data set, yielding a total of 582 antibodies (Percentage of non-detectable spots per analyzed antibodies: min = 0%, max = 51%, median = 1.3%, q10 = 0.08%, q90 = 10.2%; **Supplemental Data File**). Remaining non-detectable data points were then replaced with the greater value of either the half-minimal value across spot replicates per patient or the half-minimal value across spot replicates across patients. The data were Log2 transformed and replicates combined by averaging. Iterative row-and column-wise median centering and normalization were performed fifty times for each array type (RayBiotech or in-house) separately. The normalized data were then Z-scored along proteins, combined from both arrays, and used for analysis.

### Differential Protein Level Analysis

To identify proteins with significant changes in plasma levels we calculated permutation-corrected p-values (pcorr) for NC vs. svPPA for every protein (unpaired two-tailed t-test, 10,000 permutations). To compute false-discovery rates, we adopted a direct approach to estimate q-values (9, 25). We then rank-ordered the proteins by corrected p-value and selected the top 25 most significantly changed proteins for unsupervised clustering (CLUSTER 3.0 for OSX; M. de Hoon, Riken, Japan). The same procedure was applied to the AD comparison samples (Jaeger et al, under review).

### Differential Correlation Analysis

To gauge protein correlations, we performed a pairwise Pearson correlation between all 582 proteins for the NC or svPPA patients. Differential correlations were calculated by subtracting the respective NC correlation from the svPPA correlation. Unsupervised single linkage clustering on the correlation profiles resulted in 13 blocks of proteins with similar changes in correlation profiles (i.e. these protein correlations change alike in disease conditions). To identify those protein blocks that contribute the largest share to profile similarity, we calculated the absolute sum of correlations for each cluster and thus obtained a ranking of clusters (**Supplemental Fig. S2**).

### Biological Network Analysis

To link the differential changes in protein expression and the changes in network connectivity to biological pathways, we formed the union of all proteins significantly different between svPPA and NC (p_corr_ < 0.01, n = 33) and the proteins that are part of clusters 8 and 5 (n = 34 and n = 39) for a total of 100 unique proteins-of-interest (**Supplemental Data File**). We mapped these proteins onto known protein networks using the Genemania-app in Cytoscape 3.0.1 (www.cytoscape.org) and color-coded the protein nodes if pcorr < 0.05. Pathway data came from NCI-Nature, Reactome, and (20, 26). Physical interaction data came from Biogrid Small Scale, IREF Interact, and IREF Small Scale. To test for enrichment in biological function, we queried Gene Ontology, KEGG, and Panther databases using DAVID with the 564 unique genes representing the 582 proteins tested as background and found significant enrichment for several pathways related to cellular immunity.

### Patient Peripheral Blood Mononuclear Cells (PBMC) Acquisition

Patient and age-matched control EDTA blood was collected at the UCSF Center for Memory and Aging with UCSF IRB approval and transferred to Stanford University on ice. PBMCs were isolated by Ficoll-Paque gradient centrifugation, collected in fetal bovine serum with 20% DMSO, and slowly frozen at −80°C, where they were stored until use.

### CyTOF Immunophenotyping

Immunophenotyping was performed in the Stanford Human Immune monitoring Center. Briefly, PBMCs were thawed in warm media, washed twice, resuspended in CyFACS buffer (PBS supplemented with 2% BSA, 2 mM EDTA, and 0.1% soium azide), and viable cells were counted by Vicell. Cells were added to a V-bottom microtiter plate at 1.5 million viable cells/well and washed once by pelleting and resuspension in fresh CyFACS buffer. The cells were stained for 60 min on ice with 50uL of an antibody-polymer conjugate cocktail (**Supplemental Table S4**). All antibodies were from purified unconjugated, carrier-protein-free stocks from BD Biosciences, Biolegend, or R&D Systems. The polymer and metal isotopes were from DVS Sciences. The cells were washed twice by pelleting and resuspension in 250 uL FACS buffer. The cells were then resuspended in 100 uL PBS buffer containing 2 ug/mL Live-Dead (DOTA-maleimide (Macrocyclics) containing natural-abundance indium). After washing the cells twice by pelleting and resuspension with 250 uL PBS, the cells were resuspended in 100 uL 2% PFA in PBS and placed at 4°C overnight. The next day, the cells were pelleted and washed by resuspension in fresh PBS. The cells were resuspended in 100 uL eBiosciences permeabilization buffer (1x in PBS) and placed on ice for 45 min before washing twice with 250 uL PBS. If intracellular staining was performed, the cells were resuspended in 50 uL antibody cocktail in CyFACS for 1 hour on ice before washing twice in CyFACS. The cells were resuspended in 100 uL iridium-containing DNA intercalator (1:2000 dilution in PBS; DVS Sciences) and incubated at room temperature for 20 min. The cells were washed twice in 250 uL MilliQ water. The cells were diluted in a total volume of 700 uL in MilliQ water before injection into the CyTOF (DVS Sciences). Data analysis was performed using FlowJo v9.3 (CyTOF settings) by gating on intact cells based on the iridium isotopes from the intercalator, then on singlets by Ir191 *vs.* cell length, then on live cells (Indium-LiveDead minus population), followed by cell subset-specific gating.

### SPADE analysis

SPADE analysis was performed on PBMCs from an individual NC and svPPA patient as described in Qiu et. al using the spade.R script and Cytoscape (21, 27). Files were down-sampled to 10 percentile and clustered with a target of 100 clusters based on the values of all markers listed except DNA. Clusters were connected using a minimum spanning tree, and medians of expression values and percent totals were calculated based on up-sampled data.

### TNFα ELSIA

Mouse plasma TNFα levels were measured using the Mesoscale Mouse TNFα Ultra-sensitive kit (Mesoscale, Rockville, MD) according to the manufacturers instructions. Briefly, diluted calibrators and neat samples were loaded onto pre-coated ELISA plates and incubated for 2 hrs with vigorous shaking. Plates were washed, incubated with the detection antibody solution and then read on a Mesoscale Sector Imager after addition of 2 x Read Buffer T.

### Mouse plasma Luminex analysis

The Roberson Lab at the University of Alabama at Birmingham collected plasma from Grn+/+ and Grn-/-mice. EDTA plasma was generated by centrifugation of freshly collected blood and aliquots were stored at −80°C until use. The relative concentrations of cytokines and signaling molecules were measured in plasma samples using standard antibody-based multiplex immunoassays (Luminex) by Rules Based Medicine (Austin, TX), a fee-for-service provider.

### *Grn*^-/-^ PBMC FACS

Whole blood was collected by submandibular bleeds and clotting prevented by addition of 15 μl of 250mM EDTA. Plasma and cells were separated by centrifugation at 1000 x g for 10 min. PBMCs were subsequently collected by Ficol-Paque Plus (Sigma-Aldrich) gradient centrifugation and washed with FACS buffer (1x PBS, 1% FBS, 0.02% Sodium azide). Blocking was performed using unconjugated CD16/CD32 (FC block) for 15 min on ice and subsequently stained with fluorescently conjugated antibodies for 30 min on ice in the dark. The following antibodies were used CD3-FITC, CD4-APC-Cy7, CD8-PacBlue and CD27-Pe-Cy5.5, all from eBiosciences. Cells were washed with FACS buffer and measured on a BD LSRFortessa. Data was analyzed using FlowJo X 10.0.

